# Mutation Mechanism In DNA: Non-Hermitian Approach

**DOI:** 10.1101/2023.09.29.560200

**Authors:** Mustafa Sarısaman, Mehmet Ali Tibatan, Seval Uzunal

## Abstract

We propose a novel mutation mechanism for points and ordinary or palindromic sequences of DNA and RNA. We adopted non-Hermitian approaches based on quantum mechanics. Hermiticity is in the limelight of any physical structure with quantum character, like DNA, or RNA, as it creates quantum stability in that it yields real eigenvalues and orthonormal states. We show that, through the mutation mechanism we constructed based on non-Hermitian physics, the deterioration of the Hermitian character of the original DNA states, nucleotides, does not create a stability problem. We show that Weyl’s perturbation theory helps us determine the stability of mutated DNA or RNA. We prove that mutations made in the laboratory with conventional nucleotides using non-Hermitian physics methods are not different from mutations that occur spontaneously in nature. This result may help to reveal the quantum nature of genetic diseases in the near future and may shape the molecular approaches.

## 1. Introduction

With the development of modern science, many mysteries about the prominence and scope of DNA have been clarified by the harvest of the human mind. However, there are still many riddles to be understood regarding the magnificent structure and functions of DNA. With the understanding that DNA exhibits a quantum character, can it be possible to reveal the intriguing behaviours of DNA with quantum mechanical methods rather than simply explaining its electrochemical and,or catalytic interactions with enzymes [1] or other molecular factors? This motivation firstly provided us with a basis for understanding the quantum features in DNA, and in this way, it was possible to reveal various symmetrical structures related to DNA that are not yet known. In this way, the unitary character of DNA was studied and its extraordinary aspects were deciphered in [2], see also Refs. [3, 4, 5, 6, 7, 8, 9, 10] for the literature on some symmetrical features found in DNA. However, the necessity of understanding the mutation mechanism in DNA is one of the most important problems that await an answer. Our experience has shown that the mutation mechanism can be explained quite well, especially by non-Hermitian quantum mechanical methods, see [11, 12, 13, 14] for basics of non-Hermitian physics. It has been realized that non-Hermitian physics has a very important place in nature, especially recently, and a hectic work environment has been initiated in this direction. With this motivation, in this study, the intriguing and unknown mysteries of stable mutations are revealed by non-Hermitian physics methods by developing a perspective on the general structure of DNA and especially the mutation mechanisms in palindromic regions for the first time with quantum mechanical methods. In this way, we unravel the possibility to carry out the mutation mechanism in a controlled manner. This will enable the most important diseases of today’s world such as the treatments of genetic diseases originating from DNA and cancer to be solved. In this way, we may exhibit how the quantum mechanical properties of the DNA molecule reveal themselves on a macro-scale by showing this defective gene region effect as a disease. This might be the governing example of showing the quantum scale dynamics at the macro-scale in our daily life.

Gene sequences are a very special structure formed by nucleotides in DNA, with inversion symmetry in a double helix as a result of the DNA replication process. For instance structures like palindromic sequences, tandem repeats, polymorphic replacements, and mirror symmetry within the DNA structure provide specific information access and, or hot spots for the genetic process exhibits quantum-like properties, making these structures very interesting to examine at the quantum level. In our previous work [2], we showed that symmetrical palindromic sequences within the genetic regions have a quantum unitary structure, which concluded that vitally important information passed down from generation to generation in DNA is transmitted through this unitary feature. At this point, a very critical issue that needs to be emphasized and understood is the mutation mechanism, which happens to unveil as a result of the violation of this unitary structure. Furthermore, this mechanism is in fact quite an intricate phenomenon and can have many different causes.

In this study, we investigated the mutation mechanism in the framework of single and double-strand sequences, like palindromic ones and point mutations resulting in CAG repeats at specific gene regions of DNA. The well-known hallmark of traditional quantum mechanics is that it has a hermitian character. Therefore, the prominent aspect of any physical structure that exhibits quantum features is that it displays hermitian properties. This requires the physical system to have real eigenvalues and orthonormal eigenstates. Since it is known that the structure of DNA involves quantum properties, it is our most natural expectation that it should implicate a hermitian characteristic. Violation of hermitian structure may lead to breaking the quantum-like features, and therefore mutation incidents will not survive and will generate another mutation until a stable state is formed. The final stable state of mutated DNA will again contain a quantum unitary structure.

In this paper, we examine the mutative properties of sequences including palindromic structures and whether mutations can be carried out in a controlled manner, provided that the stability of these mutation states is ensured. First of all, we would like to emphasize that the mutation in DNA may occur either in a single strand or during replication process in both strands or in both cases together and simultaneously. However, since the second strand can be expressed by the inversion symmetry of the first strand, we will represent two clinical cases with mutation-associated genetic disorders taking place in the single DNA strand [15, 16]. We expect the mutation mechanism we have constructed to clarify many unknown situations related to DNA, and DNA transcripts (mRNA, non-coding RNAs etc.) and to offer novel approaches to understanding the nature of diseases including cancer. Within this concept, we investigated the quantum properties of mutations at E-box (CANNTG, including palindromic sequences in the e-box regions) region mutations in congenital myasthenic syndrome [15] and CGA repeats in Fragile X retardation syndrome [16]. In both cases; neuronal development is abolished and leading to mental retardation or mental developmental diseases. However, genetic stability is also distorted in these mutational systems. Genomic stability can cause the inhibition of the specific gene region process by the enzymes and, or transcriptional factors [17].

As a result of our study, we have shown that the mutation mechanism may actually be carried out in a controlled manner in the laboratory environment, if it is done with standart nucleotides known in nature. Moreover, non-Hermitian physics can provide us with a large spectrum of and difficult to control mutation environment. However, for our purposes, making mutations with conventional nucleotides is quite easy and creates a biologically stable states. This has the potential to create hope that can shed light on the solution of today’s important diseases such as DNA-related diseases and cancer.

## 2. Construction of the Formalism: Regular DNA Structure

Consider a normal DNA strand (no mutation) in any random phase that consists of a countable number of ordered distinct states. As indicated by Chargaff’s parity rules DNA shows unique symmetrical structure and frequencies within its structure [18]. However, there are irregularities which alter the genomic stability and functionality of the transcribed product. In order to construct the formalism, we stress out the nucleotide basis adenine |**A**⟩, thymine |**T**⟩, guanine |**G**⟩, cytosine |**C**⟩ and |**E**⟩ . Here |**E**⟩ represents an empty state in the absence of conventional nucleotides, see Fig. 1. We denote this set of nucleotide basis by χ⟩ such that;

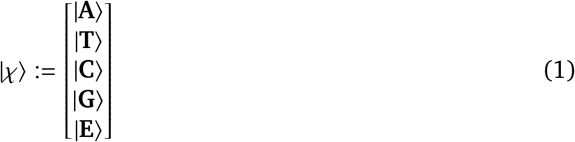

**Figure 1:**
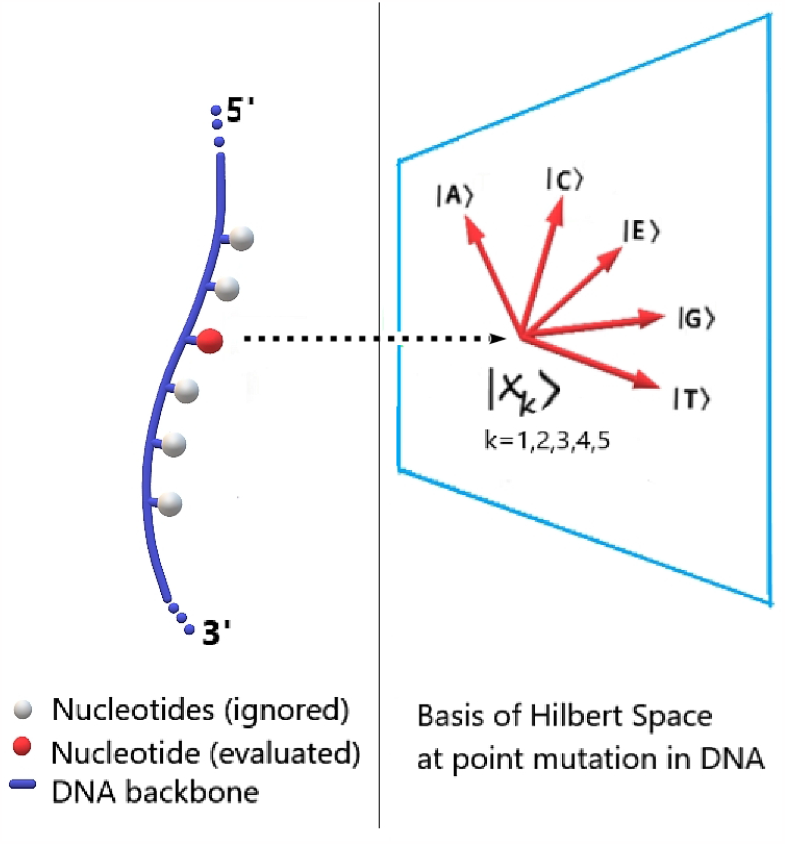
(Color online) Simplified representation of states in a single strand of DNA. Here symbol *k* denotes distinct states.

Notice that DNA replication occurs in accordance with Chargaff’s rules, which can be identified by

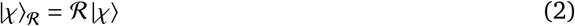

where we specify the state |χ⟩_ℛ_ to be replicated strand of DNA and the replication operator ℛ by

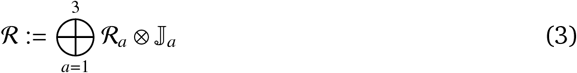

Then we realize that;

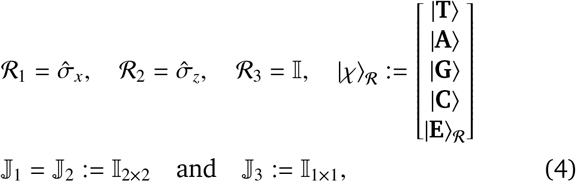

where 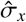 and 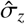 are the celebrated Pauli sigma matrices and 𝕀_*k*×*k*_ is *k*-dimensional unit matrix, see [2] for details of these conventions. In principle, we can admit |**E**⟩_ℛ_ = |**E**⟩ . At this point, we need to clarify the role of the state |**E**⟩ in our setup. The usual formation of DNA in unmutated case does not incorporate states other than the known nucleotide types. But in order to explain the null states or any molecules other than the nucleotides that may occur in the case of mutation, we had to include such a state in our construction, which we denote with E. While this can actually be a bit confusing at first, it found a complementary and quite nice place for the consistency of our formulation as the mutation did not affect the previous state.

We observe the standard relations 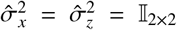 once we have repeated replication in the other strand of DNA, which is expressed by ℛ^2^ = 𝕀_5 × 5_. This is a necessary and sufficient condition for the unitarity of DNA structure.

In our previous study [2], we investigated some substantial features of palindromes in DNA in view of symmetry aspects. Inspired from the results of this study, we reveal that palindromic structures have a perfect time-reversal symmetry in the sense of quantum mechanics. One can not claim a time-reversal symmetry for non-palindromic structures. Let **T** be a time reversal operator, such that when it is acting on the state |χ⟩, the state is transformed to |χ⟩_ℛ_, **T** |χ⟩ = |χ⟩_ℛ_ . This is in fact equivalent to the replication operator ℛ. Thus, it is understood that the replication operator is endowed with the role of time-reversal operator **T** in palindromic structures. In other words, palindromes have time-symmetry. Otherwise, one can not mention about time-reversal symmetry in Chargaff’s rules in general.

### 2.1. Mutations in DNA

Based on the regular structure and functionality of DNA, by the concept of mutation, we refer to changes in the original structure of DNA throughout the article. We will not emphasize the causes of these changes in our study, mutations of interest may have been caused by any reason. As can be realized from this argument, any deformation in a typical DNA can occur in myriad ways. However, we will confine our attention to only mutations to exact known nucleotides for simplicity. Ordinary DNA is formed by four different types of nucleotides, i.e. Adenine, Thymine, Cytosine and Guanine. Nonetheless, we will not exclude the possibility of an empty state since it is one of the most commonly encountered states in mutations in general, that is why we incorporated empty state |**E**⟩ into Eq. (2).

Since mutations occur as a result of deformations in the known standard structure of DNA, they also bring many problems to be considered. First, are these mutations beneficial or harmful? Then, can these mutational states be prevented if harmful or controlled if beneficial? Are mutations stable or can stable mutations be obtained? In order to answer all these questions within the concept of mutation, it is necessary to understand how the mutation mechanism is performed. This is in fact the main subject and also the motivation of our article.

## 3. Method

Regardless of the causes and processes that create mutations, a matrix-valued operator will be defined to represent point mutations by considering only a single strand. A mutation matrix for a sequence that has multiple nucleotides will be fulfilled correspondingly by respecting the order of states. This approach will be adopted for two conjugate strands of DNA due to time-reversal symmetry of each other and then pairing matrices will be formed eventually. Similarly, palindromic sequences may be evaluated in this context.

Let ℳ be a mutation operator mapping initial state |χ⟩ to the final mutated state |χ^′^⟩,

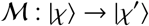

such that

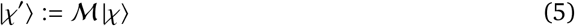

We stress out that a primed state implies a mutated state in this study. We will develop our mind map and theoretical constructs based on this configuration. In this sense, it is rather beneficial to introduce some basic concepts related to the mutation phenomenon to create a conceptual framework and consistency for the reader.

### 3.1. Conceptual Framework and Roadmap of Formation

### Genetic stability and instability

One of the most important issues that we have to overcome when dealing with the mutation phenomenon is the issue of genetic stability. Instability term refers to the distortion or inhibition of genetic regions within the DNA caused by mutations. Abnormal sequential changes in any genetic regions lead to enzymatic-transcriptional failure and end in various diseases including cancer [19]. External (radiation, toxicity, etc.) or internal factors (errors during the DNA repair or proofreading during transcription) form the mutational changes in DNA and this directly changes the protein recognition of the region.

Oncogenic regions are another factor responsible for genomic instability and these regions are directly responsible for the replication stress of DNA which drives the transcription process more susceptible to mutation-associated tumorigenesis. Naturally, activation of these regions may be responsible for DNA repair error or replication stress [20].

In any words, the genetic stability and instability terms we employ in this study indicate the functionality of specific gene regions and their information transfer throughout the generations in some cases. Not all cases have to support the mutations but DNA transcriptional system recognition is affected in any case of instability of the genetic region.

#### Quantum stability

Although genetic stability indicates the biological dimension of the mutation event, we will use the concept of “quantum stability” as the physical equivalent of this phenomenon in the quantum formalism we use in our mutation modelling. In this study, this refers to being consistent in states that are known as five nucleotides^1^ and empty or undefined molecules that are all named as states, e.g. whatever the process affects on these states the result will again be one of them, so the spectrum (the measured values of states) will be stable.

#### States and base vectors

In our setup, nucleotides are represented by state and base vectors. Each nucleotide is considered to be a state, namely generalized | χ_*k*_ ⟩, where the index *k* = 1, 2, 3, 4, 5 signifies certain nucleotides (Adenine (A), Thymine (T) or Uracil (U), Guanine (G) Cytosine (C) and Empty State (E)), and | χ_*k*_ ⟩ _ℛ_ was used for the complementary strand of double helix DNA structure.

Notice that in the case of the existence of any molecule other than the known ones, or the null point which has no nucleotide at all, a new states | *E* ⟩ and | *E* ⟩ _*R*_ was introduced, as stated in the Construction of the Problem section, section 2 of this document.

Components | χ ⟩ _*k*_ of base vectors identified at a single point are explicitly denoted as follows

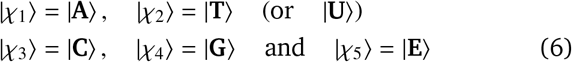

#### Sequence

This will be used for multi-states, palindromic or not, evaluated together. The same state can exist more than once in a sequence, and there is no restriction on sequence size. Each different permutation of multi-states gives rise to a new sequence.

#### Transformation

This term is used to express the task that a nucleotide is affected by an agent or process. A mutation may or may not occur as a result of a transformation. Transformation of states will be identified by the following mapping,

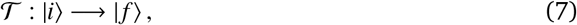

such that | *f* ⟩ = 𝒯 | *i* ⟩, where 𝒯 is the operator that makes transformation, *i* is the initial state and | *f* ⟩ is the final transformed state. For single states, this is addressed by | χ_*i*_ ⟩→ | χ _*f*_⟩, whereas, for a multi-state, transformation occurs as |χ_*i*_ χ_*i*_ … ⟩ → |χ _*f*_ χ _*f*_ … ⟩ respecting the order of the states. This is in fact denoted by the frame shift. We realize that 𝒯= ℛ in case of replication and 𝒯= ℳ in case of mutation. In replication, 𝒯 corresponds to the enzyme that makes the replication event, but in the latter case, it is the cause of the mutation event.

We remark that multi-state expression is an ordered (position-based) operation as in written sequence, not consecutive processes.

#### Mutation Matrices

In view of all these premised concepts we have introduced, we need to distinguish mutation matrices corresponding to distinct mutation events. These are specified as follows.

*Point Mutation Matrix* is a mutation matrix which transforms the initial state of a nucleotide to a final state for a single point of nucleotide base and is denoted ^*s*^ ℳ . Here, superscript *s* indicates the order of nucleotide.

*Sequence Mutation Matrix* is a unified multi-point mutation matrix for a sequence and is denoted as ℳ. It can easily be observed that ℳ := ⊕ ^*s*^ℳ.

*Pairing Matrix* is defined for the double-strand structure of DNA. This is in fact the type of mutation which occurs during the replication process. Similar to single strand case, ^*s*^𝒟_ℛ_ represents a single point pairing on the *s*^*th*^ state in sequence, whereas 𝒟 _ℛ_ is used for the whole sequence. Notice that 𝒟 _ℛ_ := ⊕ ^*s*^ 𝒟 _ℛ_. One may experience both of replication and mutation events simultaneously. In this case, the order in which the event takes place first is significant. All these configurations are clearly illustrated in a compact way in Fig. 2. In particular, the role of 𝒟 _ℛ_ is to map |χ⟩ to |χ^′^⟩ _ℛ_,

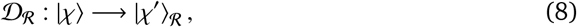

**Figure 2:**
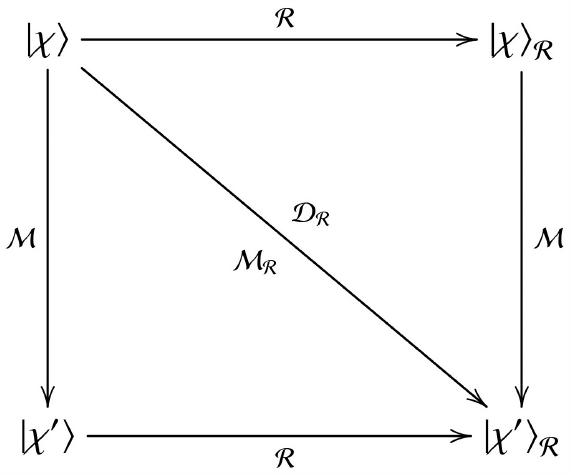
Transformation mappings show the flow of replication and mutation events of the prescribed Notice that ℳ _ℛ_ = 𝒟 _ℛ_.

**Figure 3:**
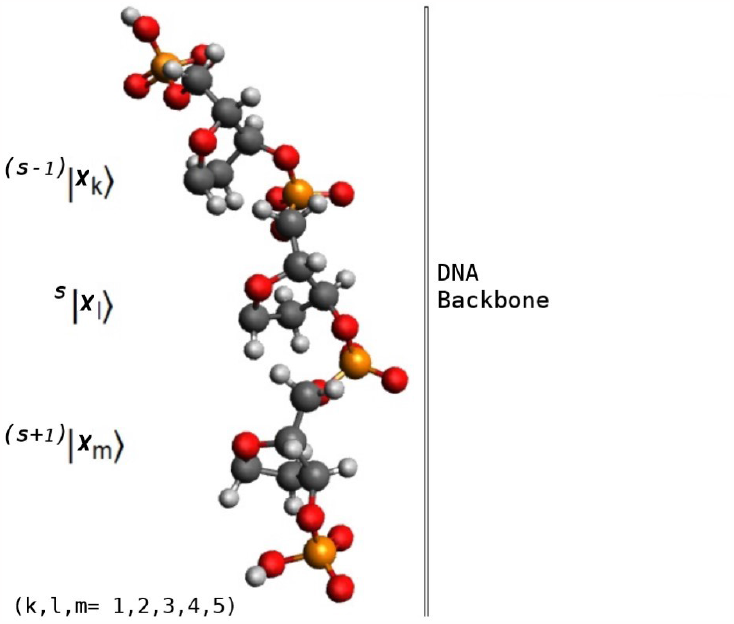
(Color Online) Representation of states on a single strand (nucleotides are removed, ball and stick model). Each state is placed as a nucleotide next to deoxyribose groups. DNA or RNA backbone; deoxyribose sugar (grey), phosphate group (orange and red). Three states shown in the figure are not codon representations, but ordinary sequences.

In Fig. 2, we realize that ℳ _ℛ_= ℛ _ℳ_ and 𝒟 _ℛ_= ℳ ℛ . The former one amounts to mutating the initial state first and then replicating. The latter implies first replication and then mutation. In principle, ℳ _ℛ_ and 𝒟 _ℛ_ have noncommutative characters, i.e. ℳ _ℛ_ ≠ 𝒟 _ℛ_, but in our case we consider a special subclass of these mappings such that ℳ _ℛ_= 𝒟 _ℛ_ without loss of generality, and they yield a commutative ordering [ ℳ, ℛ ] = 0. Therefore, 𝒟 _ℛ_= ℛ _ℳ_= ℳ _ℛ_ . This implies that there are two ways to produce . The first one is to make a mutation in the single strand and then replicate. The second one is to perform replication first, and then do a mutation. The order of the operation is insignificant. In terms of states, this special two-way process will be ( |χ^′^⟩ ) _ℛ_ = ( |χ⟩_ℛ_ )^′^. In the realm of biology, the existence of maps and are the origins of diseases as mutated sequences become erroneous gene regions in the replication process which also affects the genomic stability through the DNA ligation [21]. Thus, we may represent the defective mutations as related to the DNA replication process (mutation on lagging or leading strand of DNA) depicted in Fig. 2.

### *3.2*. *Framework of Mutation Mechanism*

#### Single strand evaluation for DNA and RNA

In accordance with the discussion stated in Fig. 2, although the mutation process may arise on a single strand or a double strand, it would not be inaccurate to consider both as essentially equivalent. Hence, evaluating only singlestrand mutation appears to be satisfactory for our purposes. In practice, although there is no limit on the number of nucleotides that are mutated, mutations in lengthy sequences require voluminous matrices difficult to work with. That is why we focus our initial attention to point mutations on a single strand but also consider mutations at multiple points of any sequences like palindromic structures as well.

We remark that mutations may lead to innumerable different states from the initial state. As stated before, we will confine the mutated states to the already-known nucleotides. Thus, if

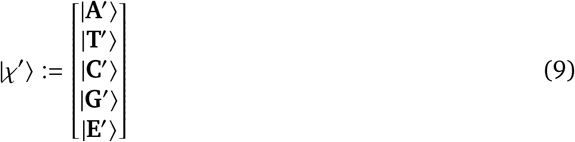

then, components of the mutated states could only belong to a closed set consisting of {|**A**⟩, |**T**⟩, | **C**⟩, | **G**⟩ and |**E**⟩, i.e., 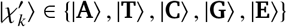 . This is possible, only if all |ϕ^′^⟩ ∈|ϕ⟩, where ϕ := **A, T, C, G** and **E**. In the rest of the paper, we will stick to this commitment.

Notice that every point transformation may not be counted as a mutation of interest. If a state (nucleotide) is transformed into itself (self-transformation), as expected, it is not a mutation. For every *k*, ℓ = 1, 2, 3, 4, 5 and *k* = ℓ

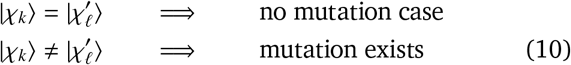

More clearly, if we look the other way, for *k* ≠ ℓ, the mapping 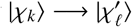 indicates the desired mutation process.

Every point may initially have any of five states and be transformed into any of them at the final stage, so there are 25 mutually distinct cases.

The point mutation matrix covers all possibilities. Hence, its entries are described by

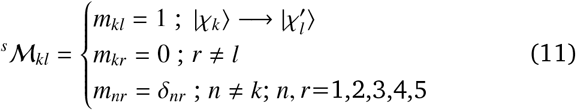

where *s* stands for the order of points in a sequence. In case no mutation exists, ^*s*^ ℳ is a unit matrix, i.e. ^*s*^ℳ = 𝕀 or in components *m*_*kl*_ = δ_*kl*_, with *k, l* = 1, 2, 3, 4, 5.

In this study, all cases are assumed to be equally likely. The existence of some cases of unequal probabilities should be investigated and we expect that the fundamentals of this formalism are still held to be true (valid) by a weighted matrix. For a sequence, all point matrices of the sequence are unified as ℳ. Superscript *s* is used to identify points, and sub-indices *k, l* are dropped from ^*s*^ℳ_*kl*_ for simplicity

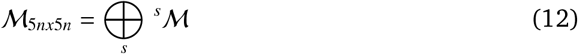

Here, *n* is the number of nucleotides in the sequence and ⊕ is the direct-sum symbol. The direct sum is over all individual points. The sequence mutation matrix for a single strand, more clearly, is as follows :

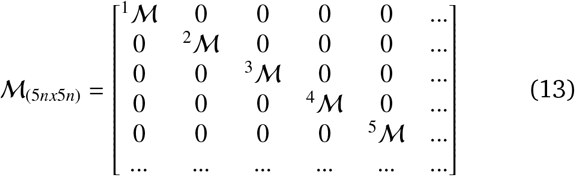

#### DNA Helix Structure, Double strand evaluation

Now, we consider a mutation case in the double strand of DNA helix structure, which takes place after replication process. Since idea is the same as before, we follow the similar approach. Notice that if there is no mutation, all pairs on complementary strand obey Chargaff’s rules. Two state vectors are paired accordingly and denoted by the mapping : 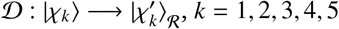 and the point pairing matrix, is the unit matrix, then ^*s*^𝒟 = 𝕀

Pair mutation matrix was defined as:

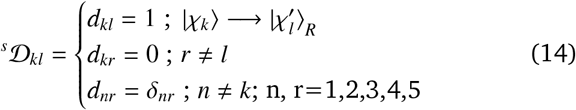

A mutation matrix of double-strand helix structured sequence, whether palindromic or ordinary, was defined as similar to a sequence mutation matrix of a single strand as follows:

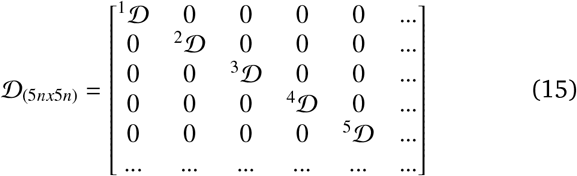

### 3.3. Quantum Stability Evaluation

One of the important points is whether the resulting states are stable or not after the mutation process is carried out. This leads to the need to perform quantum stability analysis. Quantum stability is evaluated based on Weyl’s Perturbation theorem in the context of singular value spectra [12].

THEOREM 1 (Stability of singular value spectra) *For two arbitrary nxn matrices 𝒰 and 𝒫, denoting for the j*^*th*^ *largest singular value of 𝒰 and that of 𝒰* + 𝒫 *as* γ _*j*_ *and* 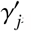, *respectively, we have*

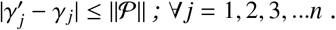

In this theorem, the matrix 𝒫 denotes the perturbation of the non-Hermitian matrix 𝒰 + 𝒫 such that the matrix 𝒰 is hermitian. Although this theorem holds for all 𝒰, 𝒫 ∈ ℂ^*n*×*n*^, in our case 𝒰, 𝒫 ∈ ℝ ^×*n*^ since our mutated states involve only known nucleotides.

THEOREM 2 (Stability against general perturbations) *For an arbitrary (nxn) Hermitian matrix* 𝒰 *and an arbitrary (nxn) matrix 𝒫, denoting the largest eigenvalue in* Λ(U) *as* λ _*j*_, *we have*

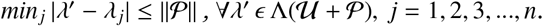

This theorem implies that the spectra of Hermitian matrices are also robust against non-Hermitian perturbations. The evaluation has also been done by using real case data, see Fig. 5.

**Figure 4:**
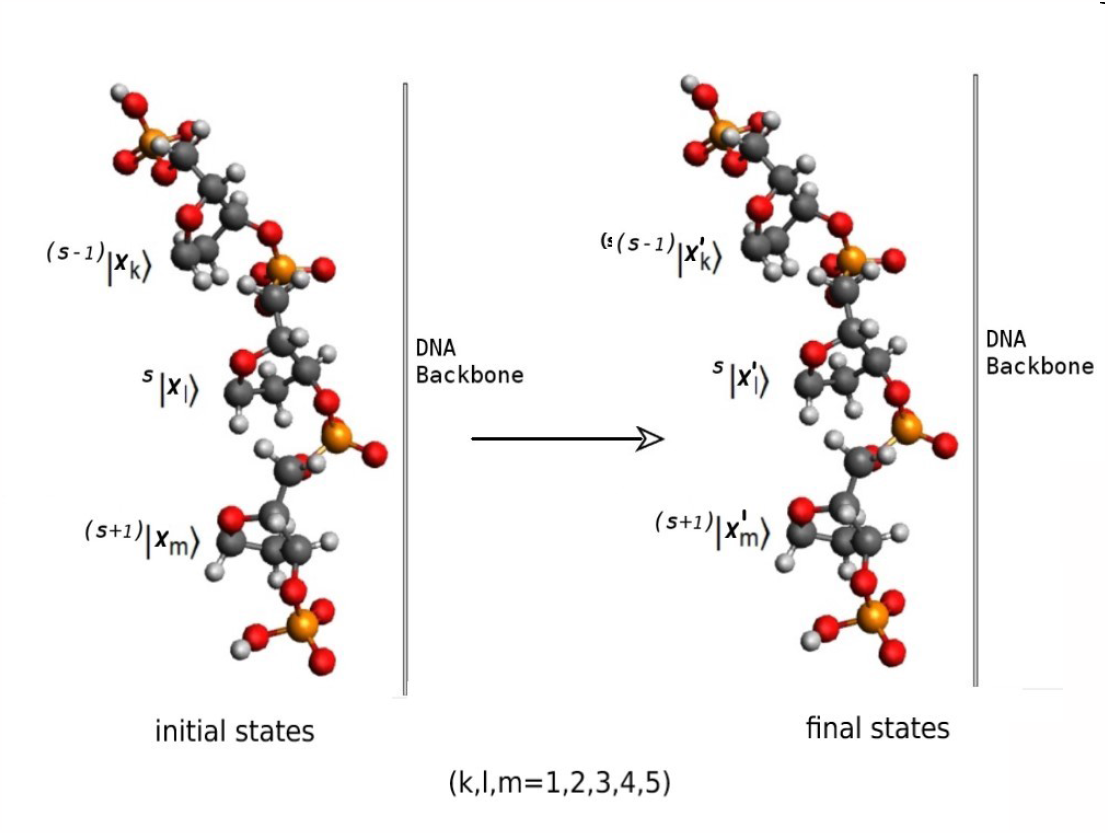
(Color Online) Transformed (i.e. mutated) states on a single strand of DNA (nucleotides removed, ball and stick model). Every state has initial and final (shown as primed) values based on causes and processes in between.

**Figure 5:**
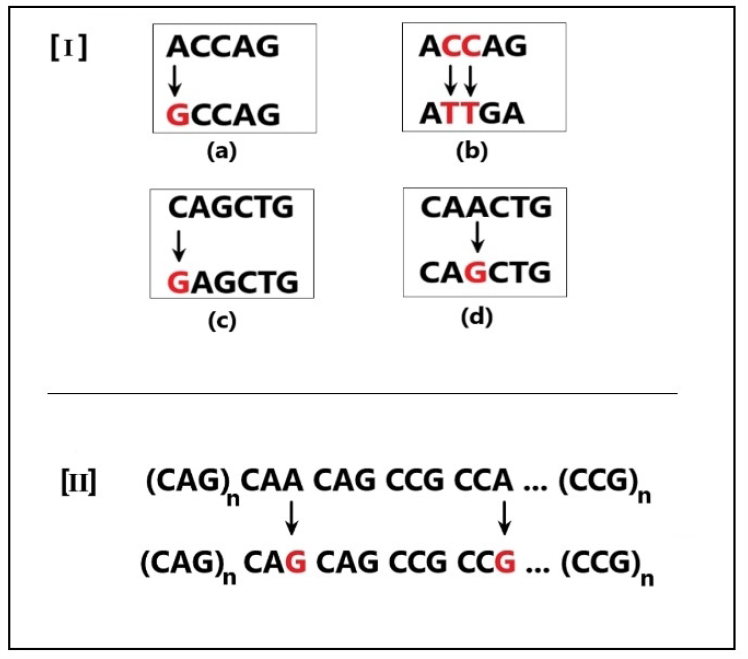
[I] Point mutations in RAPSN gene promoter region ; [II] Nucleotide replacement cause CAG repeats directly associated with genomic stability and mutational switchover of the DNA transcriptional system.

The real cases provided in Fig. 5 are related to the E-box mutation in the promotor region of the RAPSN gene in myasthenic syndrome (I) and the CAG repeats which are associated with Fragile X syndrome (II). Adenine replacement with Guanine leads to CAG repeats extension in the region and causes genetic instability of the Htt gene which is responsible for molecular transport and signalling cascade. Similar nucleotide replacement during replication in the E-box regions alters the binding affinity of related proteins and results in facial malformations in some populations [15, 16].

### 3.4. Evaluation of Quantum Stability on real case data

Oncogenic regions are another factor responsible for genomic instability **Case [I]:**This case is provided in [15].The four sequences shown in Fig. 5 explicitly, were investigated. These four cases in fact reflect the E-box mutations in the *RAPSN* promoter region with congenital myasthenic syndrome. We analyze them correspondingly, and compute the stability analysis based the above theorems.

#### First data:(a)

There is one point mutation in the genetic structure of families having *RAPSN* mutations[15].

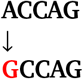

While |χ_1_⟩→ | χ_1_⟩ had been expected, meaning that no mutation exists, in fact | χ_1_⟩ →| χ_4_⟩ occurs, i.e. | **A**⟩ →| **G**⟩ mutation eventualizes.

In this situation, no mutation case is represented with 𝒰= 𝕀, and the process causing the mutation is assumed to be a perturbation noted as 𝒫, then 𝒰+ 𝒫 = ℳ and hence one obtains 𝒫= ℳ − 𝒰

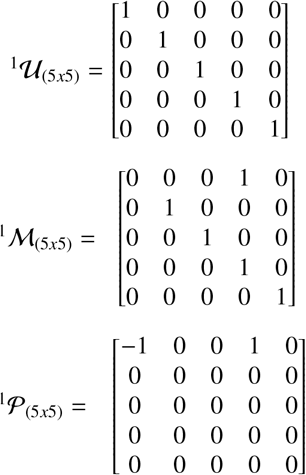

Sequence related matrices are constructed as follows once point mutation matrices are obtained. Notice that each element of the matrices ℳ _(25*x*25)_ and 𝒫 _(25*x*25)_ represents (5*x*5) matrices.

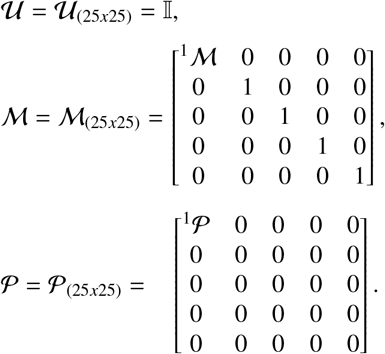

Here, ℳ and 𝒫are not Hermitian matrices (in our case, real valued non-symmetric matrices). According to Weyl’s perturbation theorem, stability of states in terms of the singular value spectra is guaranteed by the condition 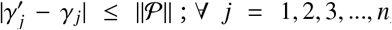, if λ _*j*_ and 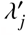 are the eigenvalues, and γ _*j*_ and 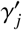 are the singular values of the operators 𝒰^*T*^ 𝒰 and ℳ^*T*^ ℳ respectively. Therefore,

By definition provided in [12, 22], one gets the following

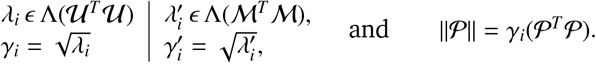

Only different eigenvalues of spectrum were stated, in accordance to roster form, with their multiplicity as (λ_*i*_;algebraic multiplicity). For example, (3;25) means that 3 is repeated 25 times in the spectrum of the matrix; where 3 is the eigenvalue and 25 is an algebraic multiplicity of 3.

Computing the corresponding quantities yields the following values:

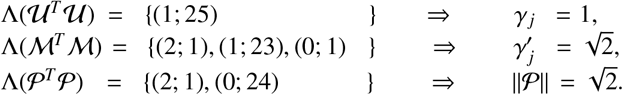

Hence, it is easy to see that the necessary stability condition 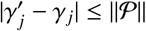 is satisfied by 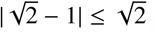.

Thus, we conclude that this natural mutation is stable as is the case. Our non-hermitian approach verifies this naturally occurring mutation process.

#### Second data:(b)

Now, we investigate the two-point mutations given in [15] as follows

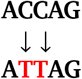

Notice that both point mutations arise as | χ_3_⟩ → | χ_2_⟩, (e.g. | **C**⟩→ | **T**⟩ ) and obviously they take place in the second and third positions in the sequence, then the point mutation matrices take the following forms:

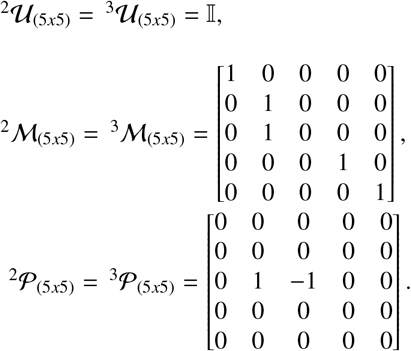

Therefore, the mutation matrices for the sequence turn out to be:

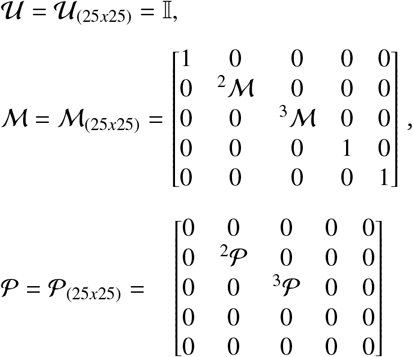

Having (25*x*25) sized matrices, namely 𝒰, ℳ and 𝒫, gets our problem a little sophisticated. But fortunately, because of entries consisting of real positive integer values, we pass through the edge of difficulty in calculations.

For both the sequence, the mutation and perturbation matrices are block diagonal, their spectra can be obtained as a union of each diagonal matrices spectrum, therefore a singular value which is based on the maximum value of the spectrum of the mutation matrix, and the norm of the perturbation matrix were calculated as follows:

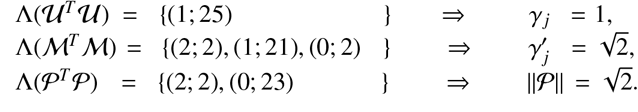

Hence, again stability condition 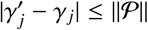 is provided by 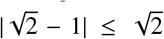 as expected. This result confirms the stability analysis of naturally occurring mutation that we presented in our approach.

#### Third data:(c)

Next, we study the stability analysis of point mutation given in congenital myasthenic syndrome study [15],

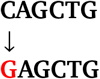

It is obvious that the mutation appears to be | χ_3_⟩ → | χ_4_⟩ and it is located in the first position in the sequence, then the point mutation related-matrices turn out to be as follows,

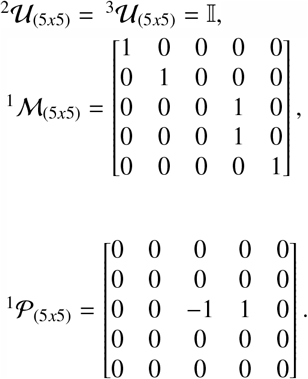

Meanwhile, for the sequence, the related sizable matrices are computed to yield

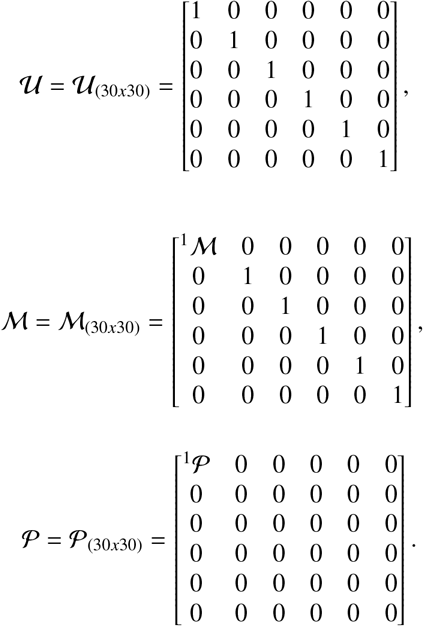

Notice that each element of the sequence mutation matrices above, represents (5*x*5) matrices. Thus, total mutation matrices build up by (5×5) matrices and have size (30×30) for this sequence. After some lengthy computations, one attains the following

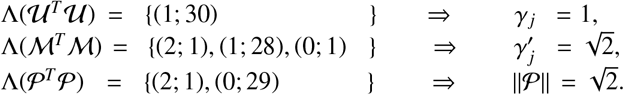

Hence, the stability condition 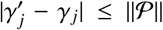 is satisfied by 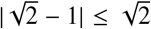. Once again, our approach supports the stability of the mutated states.

#### Fourth data:(d)

Our last nucleotide sequence example [15] comes as one-point mutation, which occurs as

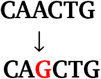

The corresponding mutation in our formalism comes out to be | χ_1_⟩→ | χ_4_⟩, which takes place in the first position in the sequence. Therefore, point mutation related-matrices will be

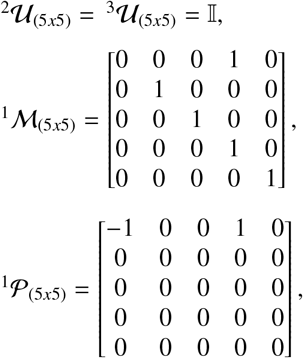

Thus, in this specific case, the relevant sequence matrices show up as follows

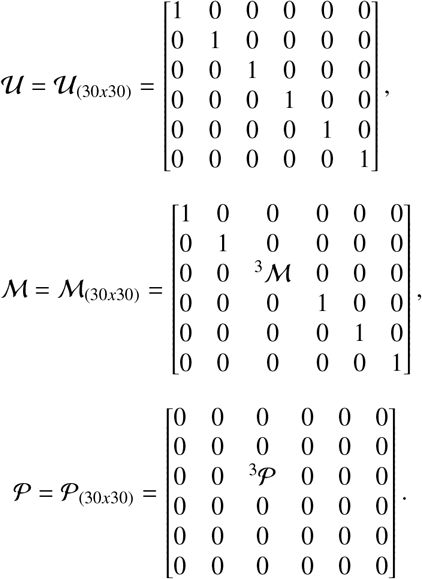

We,again, remark that each element of the sequence related-matrices above represents (5*x*5) matrices, whereas corresponding sequence related matrices consist of (30*x*30) matrices. Thus, the singular values of the respective matrices will be

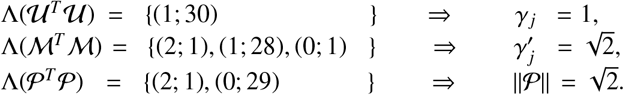

Once again, the stability condition 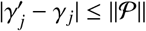 is met by 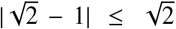, verifying that this mutation process is compatible with our formalism.

**Case II:** This case is grounded on the DNA sequence and **CAG** size to mutation frequencies of intermediate alleles for Huntington disease [16]. See the lower panel in Fig. 5 and Fig. 6 displays the sipmplified description of the mutated state.

**Figure 6:**
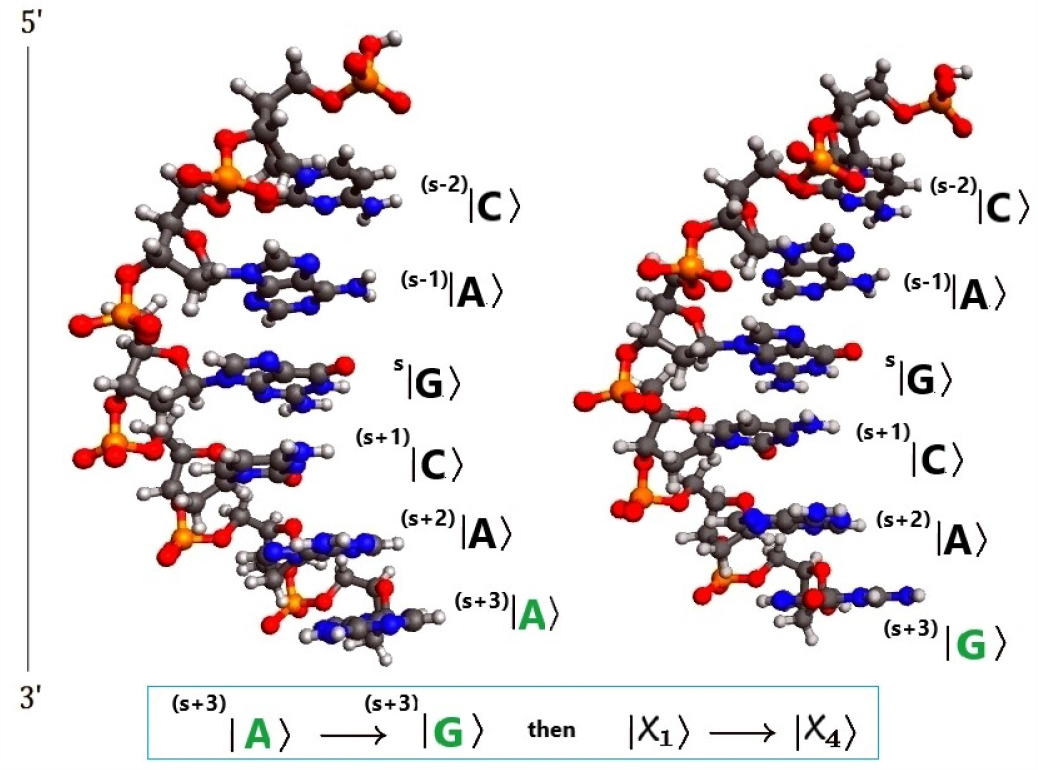
(Color Online) Ball and stick representation of the point mutation causes CAG repeats which leads to genetic instability (*s* + 3)^*th*^ [16]. Notice that |**A**⟩ is mutated to |**G**⟩.

The corresponding mutations occur as a result of mapping |χ_1_⟩ → |χ_4_⟩, (e.g. |**A**⟩ → |**G**⟩) and they are positioned in the 3^rd^ and 12^th^ places in the sequence, and 𝒰_60*x*60_ = 𝕀,

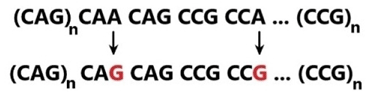

was evaluated.

Point mutation and perturbation matrices are figured out to be :

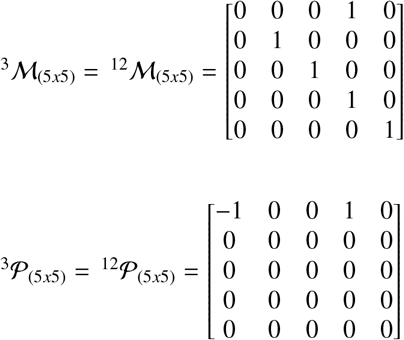

Hence, sequence mutation and perturbation matrices are constructed as follows:

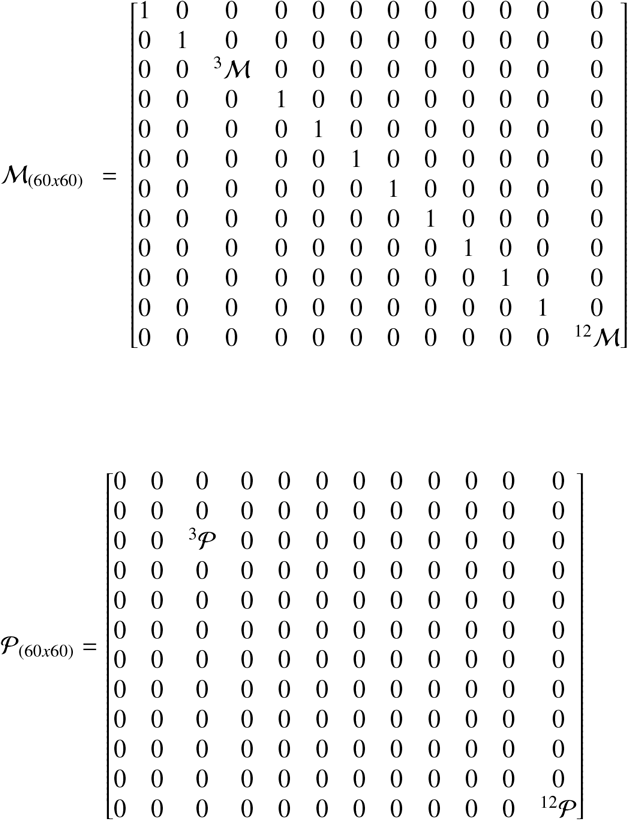

Again, both sequence mutation and perturbation matrices are block diagonal forms and their spectra can be obtained as the union of each diagonal matrice’s spectrum.

One finds out the computed values as follows

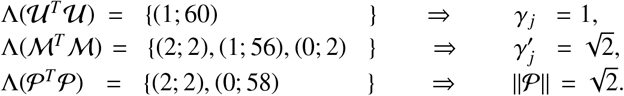

As a result of these findings, stability condition 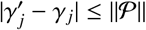 is satisfied as seen by 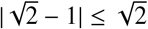.

All these results we have obtained from real data show the accuracy of the formalism we have constructed on the basis of nonhermitian physics. As can be seen, all mutation matrices that comply with our formalism will always produce stable mutations, since they consist of known nucleotides. We express this clearly with the following theorem.

THEOREM 3 (Stability of Mutations to the Conventional Nucleotides) *Let ℳand 𝒟 be the matrix valued-mutation operators of sequences for conventional nucleotides* ***A, T, C*** *and* ***G*** *in single and double strands of DNA, respectively. Then, the corresponding mutation operators ℳand 𝒟 are always stable, as proven by the Weyl’s perturbation theorem*.

This finding, which we revealed as a result of the approach we put forward in our study, has very important consequences. Once natural mutations to conventional nucleotides comes out to be stable, if viewed backwards, mutations to conventional nucleotides in laboratory environment are also stable. Both cases are in fact equivalent in this sense, there is no difference from each other. This result is very important in terms of controlling mutations. Instead of waiting for mutations to occur naturally, we gain the ability to obtain mutations in the light of the desired results in the laboratory environment. In this sense, our work opens the door to controlling DNA using non-Hermitian physics. It is clear that mutation processes other than conventional nucleotides are also worth examining and will produce more detailed results. We will not discuss these in this study due to their indepth inclusions.

## 4. Conclusion

Mutational changes within the DNA lead to genetic disorders and these disorders can be inherited by genomic information systems for generations. This informational transfer system by DNA is responsible for differences between races, genetic deformations, or genetic disorders and transferring all these changes throughout the generations. However, despite the erroneous information transferring through the generations, DNA has some conservative behaviour in the manner of providing hotspots for enzymatic recognition and/or specific protein synthesis, such as palindromic sequences [1], E-box [23], **TATA**-box or **CAG** repeats [20]. Information conservation and transferring this data through generations with errors should be investigated to reveal the origin of molecular recognition and the transcriptional process of DNA. Protective behaviours of genetic information might be answered by the quantum properties of the nucleotide. Information transfer by DNA covers viral pathogenesis, inflammation response, DNA repair mechanism signal to the proteins, transcription, translation, and DNA replication process in the cell. This complex machinery and information transfer process cannot be explained solely by biological or chemical principles. To understand the nature of DNA itself, we develop quantum-based explanations. The instability of the information in DNA is directly associated with many genetic disorders, hereditary cancer, sporadic cancers, and many other physiological abnormalities. Mutations interfering with the information process of the DNA have a complicated mechanism and use the hot spots or sensitive areas of the DNA, such as palindromic sequences, mirror-symmetrical regions, or CpG islands, which are intricated with the epigenetic mechanism [24]. By revealing the quantum mechanical behaviours of DNA sequences, we can also take an important step toward better understanding the regulatory relationship between the epigenetic system and the transcriptional system. If we consider the fact that the molecular structure and information process complexity of the DNA is complex, we may begin to see the need for understanding the quantum behaviour of the nucleotides to explain the information process and information transfer ability of our genomic system, which is ruled by DNA. We assumed all the reasons (causes) leading to the mutation as a perturbation and defined the state | **E**⟩ to identify all unpleasant situations like being empty or any molecule other than known nucleotides. These assumptions need to be investigated in detail. In our study, we focused on the mutational mechanism by using the quantum properties of normal and abnormal sequences and their functionality. If we consider the fact that mRNA is the key factor in carrying genetic encryption to be converted into meaningful protein structures, our quantum-based formalism might apply to RNA and other transcript products of DNA and even non-coding RNAs. We saw that even mutations have quantum stability in a state space that has limited definite discrete states, but this doesn’t disregard the requirement of the proper enzymes to perform the expected function. The quantum stability has no contradiction with the co-process of enzymes and DNA sequence; in fact, it points to the necessity. In this study, we formalize how a signal-coding and decoding molecule (DNA) causes clinical outcomes or how the molecular structures show their effect in the macroscale world.

Artificial genetic mutations in the laboratory are commonly used to reveal or treat the oncogenic gene regions and they have no differences from the natural mutations in the manner of mechanisms. For instance, controlled Ras mutations presented us oncogenic mechanism of some gene regions [25, 26]. Even their transcribed copy number requirements might show differences or protein levels may differ in vitro and in vivo [25] mechanism might be same. Thus, natural or artificial mutative structures on the DNA represent the same effect. This might help to understand the DNA structure and DNA resistance to mutation mechanisms at the quantum level by using the non-Hermitian behaviour of nucleic bases. In this way, we may provide a novel approach to better understand how these microscale molecules change the conditions at the macrolevel. Moreover, we use two different gene mutation situations, and both show their effects in different ways. However, E-box mutations on promotor regions of DNA or **CAG** repeat problems have the same non-Hermitian behaviour. Quantum states we formalize are the proof that the same type of mutations reveal themselves as two different disease mechanisms because they work the same at the non-Hermitian quantum scale. In both situations, there is a mutation that causes genetically unstable regions by their non-Hermitian behaviour on DNA, causing developmental delay.

## 5. Remarks

### Concluding Remarks

All authors contributed equally to the whole work including calculations, analysis and manuscript typing.

### Competing Interests

The authors declare no competing interests.

## Footnotes

1 These are in fact Adenine, Thymine, Guanine and Cytosine in DNA, but Urascil in RNA.

## Notes

### Competing Interest Statement

The authors have declared no competing interest.

## References

[1] M. Allevato, E. Bolotin, M. Grossman, D. Mane-Padros, F. M Sladek, and E. Martinez. “Sequence-specific DNA binding by MYC/MAX to low-affinity non-E-box motifs., volume 12. PLoS One, 2017.

[2] MA. Tibatan and M. Sarisaman. Unitary structure of palindromes in dna. Biosystems, 211(104565):1–8, 2022.

[3] JM. Girault and S. Ménigot. Palindromic vectors, symmetropy and symmentropy as symmetry descriptors of binary data. Entropy, 24 (82), 2022.

[4] C. Cattani. Fractals and hidden symmetries in dna. Mathematical problems in engineering, 2010.

[5] S. Petoukhov, E. Petukhova, and V. Svirin. Symmetries of dna alphabets and quantum informational formalisms. Symmetry: Culture and Science, 30(2):161–179, 2019.

[6] C. Cattani. Complexity and symmetries in dna sequences. Biological Knowledge Discovery Handbook, pages 93–127, 2013.

[7] S. Shporer, B. Chor, S. Rosset, and D. Horn. Inversion symmetry of dna k-mer counts: validity and deviations. BMC genomics, 17(1): 1–13, 2016.

[8] VI. Puller and SV. Rotkin. Helicity and broken symmetry of dna-nanotube hybrids. Europhysics Letters, 77(2):27006, 2007.

[9] MN. O’Brien, M. Girard, H.Lin, Millan, MO. Cruz, B. Lee, and C.A. Mirkin. Exploring the zone of anisotropy and broken symmetries in dna-mediated nanoparticle crystallization. Proceedings of the National Academy of Sciences, 113(38):10485–10490, 2016.

[10] T. Pal, R. Modak, and BP. Mandal. Dna unzipping as pt-symmetry breaking transition. 1 2023. URL https://arxiv.org/abs/2212.14394.

[11] N. Moiseyev. Non-hermitian quantum mechanics. 2011.

[12] Y. Ashida, Z. Gong, and M. Ueda. Non-hermitian physics. Advances in Physics, 69(3):249–435, 2020.

[13] CM. Bender. Making sense of non-hermitian hamiltonians. Reports on Progress in Physics, 70(6):947, 2007.

[14] R. El-Ganainy, KG. Makris, M. Khajavikhan, ZH. Musslimani, S. Rotter, and DN. Christodoulides. Non-hermitian physics and pt symmetry. Nature Physics, 14(1):11–19, 2018.

[15] K. Ohno, M. Sadeh, I. Blattand JM. Brengman, and AG. Engel. E-box mutation in the rapsn promoter region in eight cases with congenital myasthenic syndrome. Human Molecular Genetics, 12(7): 739–748, 2003.

[16] SS. Chong, E. Almqvist, H. Telenius, L. LaTray, K. Nichol, B. Bourdelat-Parks, YP. Goldberg, BR. Haddad, F. Richards, D. Sillence, C. Greenberg, E. Ives, G. Van den Engh, MR. Hughes, and MR. Hayden. Contribution of dna sequence and cag size to mutation frequencies of intermediate alleles for huntington disease: evidence from single sperm analyses. Human Molecular Genetics, 6 (2):301–309, 2 1997.

[17] WC. Chou, AL. Hawkins, JF. Barrett, CA. Griffin, and CV. Dang. Arsenic inhibation of telomerase transcription leads to genetic instability. Journal of Clinic Investigation, 108(10):1541–1547, 2001.

[18] M. Rosandić, I. Vlahović, M. Glunčić, and V. Paar. Trinucleotide’s quadruplet symmetries and natural symmetry law of dna creation ensuring chargaff’s second parity rule. Journal of Biomelecular Structure and Dynamics, 34(7):1383–1394, 2016.

[19] S. Negrini, VG. Gorgoulis, and TD. Halazonetis. Genomic instability, and evolving hallmark of cancer. Molecular Cell Biology, 11(3): 220–228, March 2010.

[20] P. Kotsantis, E. Petermann, and SJ. Boulton. Mechanisms of oncogene-induced replication stress: Jigsaw falling into place. 8 (5):537–555, 2018.

[21] JS. Williams, PP. Tumbale, ME. Arana, JA. Rana, RS. Williams, and TA. Kunkel. High-fidelity dna ligation enforces accurate okazaki fragment maturation during dna replication . ScienceDirect Journal, 12(1):4824, 2021.

[22] K. Kanatani. Linear Algebra for Pattern Processing. Springer Nature, 2021.

[23] H. Fujigasaki, JJ. Martin, PP. De Deyn, A. Camuzat, D. Deffond, G. Stevanin, B. Dermaut, C. Van Broeckhoven, A. Dürr, and A. Brice. Cag repeat expansion in the tata box-binding protein gene causes autosomal dominant cerebellar ataxia. Brain, 124(10):1939–1947, 2001.

[24] JW. Hyeon, AH. Kim, and H. Yano. Epigenetic regulation in huntington’s disease. ScienceDirect Journal, 148:105074, 2021.

[25] VY. Hua, WK. Wang, and PH. Duesberg. Dominant transformation by mutated human ras genes in vitro requires more than 100 times higher expression than is observed in cancers. Proceedings of the National Academy of Sciences, 94(18):9614–9619, 1997.

[26] Ö.Le Roux, NLK. Pershing, E. Kaltenbrun, NJ. Newman, JI. Everitt, E. Baldelli, M. Pierobon, EF. Petricoin, and CM. M Counter. Genetically manipulating endogenous kras levels and oncogenic mutations in vivo influences tissue patterning of murine tumorigenesis. (e75715), 9 2022.

